# Multimodal reference brain atlas of adult *Danionella cerebrum*

**DOI:** 10.64898/2026.03.09.710483

**Authors:** Mykola Kadobianskyi, Jörg Henninger, Daniil Markov, Antonia Groneberg, Johannes Veith, Marc Renz, Kutay Deniz Atabay, Peter W. Reddien, Leonard Maler, Benjamin Judkewitz

## Abstract

Mechanistic accounts of brain function require a common coordinate system in which structural, molecular and functional data can be integrated and compared across individuals. The teleost genus Danionella is unique among vertebrates in retaining lifelong transparency, allowing non-invasive, cellular-resolution functional imaging across the entire adult brain. A reference atlas in this model would therefore provide a strong foundation for causal and comparative circuit studies. Here we present an integrated anatomical, molecular and functional reference brain for adult *Danionella cerebrum* as a standardised atlas resource. Using a transgenic nuclear fluorescence marker, whole-mount tissue clearing and high-resolution two-photon microscopy, we generated an average reference brain from 21 adult fish to create a common coordinate system. Whole-mount in situ hybridisation for 29 neuronal markers, complemented by tract annotation from structural imaging and tracer injections, enabled us to segment 203 neuroanatomical regions. We found pronounced sex differences in telencephalic, cerebellar and hindbrain nuclei, revealing sexually dimorphic organisation across multiple brain regions. All data and segmentations are made openly accessible, providing a community resource for studies of circuit function, molecular makeup and sexual dimorphism in an optically accessible adult vertebrate brain.

## Introduction

Reference brain atlases are essential infrastructure for modern neuroscience, enabling integration of structural, molecular, and functional datasets across individuals and studies within a standardized coordinate system (Ding et al., 2016; Wang et al., 2020).

In zebrafish, atlases such as mapzebrain (Kunst et al., 2019; Shainer et al., 2023) and Z-Brain (Randlett et al., 2015), support mapping whole-brain activity to annotated brain regions, molecular markers, transgenic lines, and connectomes at cellular resolution. More recently, the Adult Zebrafish Brain Atlas (AZBA) extended the volumetric anatomical coverage to the adult zebrafish, providing a segmented volumetric reference based on the updated seminal zebrafish neuroanatomical atlas (Kenney et al., 2021; Wullimann et al., 1996). However, tissue size, opacity and the ossified skull restrict optical access to the adult zebrafish brain (but see Chow et al., 2020), precluding brain-wide *in vivo* imaging and highlighting the need for vertebrate models that retain whole-brain optical accessibility into adulthood.

To enable whole-brain imaging in an adult vertebrate, another danionin species, *Danionella cerebrum,* has been recently introduced as a model to neuroscience (Britz et al., 2021; Penalva et al., 2018; Schulze et al., 2018). *Danionella* species combine small size with lifelong transparency, permitting whole-brain optical access and *in vivo* imaging across all developmental stages due to the absence of skull ossification above the brain (Akbari et al., 2024; Britz et al., 2009; Conway et al., 2021; Tatarsky et al., 2025). These properties, combined with rich social and acoustic repertoire, make *Danionella* uniquely suited for systems-level studies of vertebrate brain function (Atabay et al., 2026; Demarchi et al., 2025; Fouke et al., 2025; Hoffmann et al., 2023; Lam, 2022; Lee & Briggman, 2023; Penalva-Tena et al., 2025; Rajan et al., 2022; Ruetten et al., 2025; Tatarsky et al., 2022; Veith et al., 2024; Yu et al., 2025; Zada et al., 2024, 2026). This creates a need for a standardised volumetric neuroanatomical resource that combines molecular identification of cell types, functional activity recordings, and anatomical segmentation, and that enables cross-modality and cross-study dataset integration.

To address this, here we present a multimodal adult brain atlas for *Danionella cerebrum* that integrates whole-brain anatomical, molecular and functional recordings. We first constructed a reference brain template from cleared whole-mount preparations imaged at cellular resolution. We then used whole-mount hybridisation chain reaction (HCR 3.0, Choi et al., 2018) to acquire molecular marker datasets. We registered 29 neuronal markers to the reference brain and combined these data with neuronal tracing and confocal reflectance-based tract imaging. These datasets enabled us to annotate over 200 neuroanatomical regions following established zebrafish nomenclature. Further mapping of whole-brain visually- and auditory-evoked activity enabled the integration of anatomical, molecular and functional datasets at cellular resolution in an adult vertebrate brain. Finally, this framework also revealed gross morphological differences between female and male *Danionella cerebrum*, providing a foundation for future studies of brain sexual dimorphism in this species. To facilitate broad use of this resource, we provide a versioned, structured, and expandable data repository, as well as a publicly available *Danionella cerebrum* brain viewer (atlas.danionella.org) that allows interactive exploration of all anatomical, molecular, and functional datasets.

## Results

In order to co-register all imaging modalities into a common coordinate system, we first generated a three-dimensional reference brain of *Danionella cerebrum,* which served as the target for registration of all subsequent datasets (Fig. 1). Given the small size and fragility of the *Danionella cerebrum* brains, this required adapting a whole-mount optical clearing protocol that allowed us to image entire intact heads, thereby minimising tissue distortion (see Methods). We then extended this clearing approach to molecular labellings and developed a protocol compatible with HCR 3.0 chemistry, an enzyme-free RNA signal amplification technique that showed the highest promise for whole-mount RNA labelling (Fig. 2; Choi et al., 2018; Lovett-Barron et al., 2020; Shainer et al., 2023). This allowed us to visualise neuronal marker distributions across the entire brain and to identify and delineate all brain regions from the Wullimann and AZBA zebrafish neuroanatomical atlases (Fig. 3).

**Figure 1.**
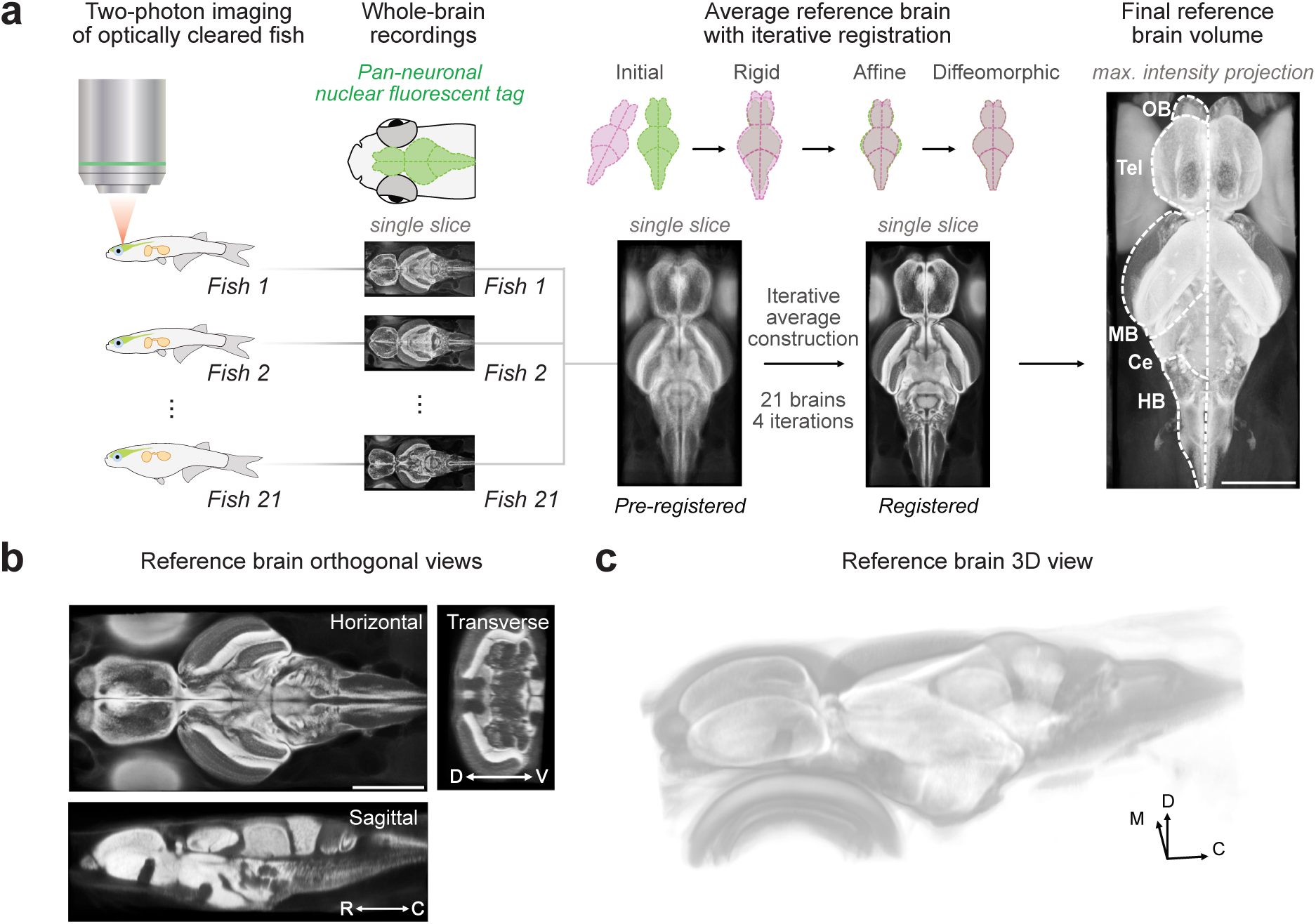
Overview of reference brain creation. (a) An overview of the pipeline: two-photon recordings of whole brains of transgenic fish expressing nuclear GCaMP6s under a pan-neuronal promoter (Tg(elavl3:H2B-GCaMP6s)), followed by an iterative average brain template construction using ANTsPy. Resulting final reference volume on the right with major divisions of the brain annotated. OB, olfactory bulbs, Tel, telencephalon, MB, midbrain, Ce, cerebellum, HB, hindbrain (b) Orthogonal projections of the reference brain middle slices in the horizontal, transverse, and sagittal views. D-V, dorsal-ventral, R-C, rostral-caudal (c) 3D rendering of the reference brain created in Fiji using the Volume Viewer plugin. M, medial, D, dorsal, C, caudal. All scale bars, 500 µm.

**Figure 2.**
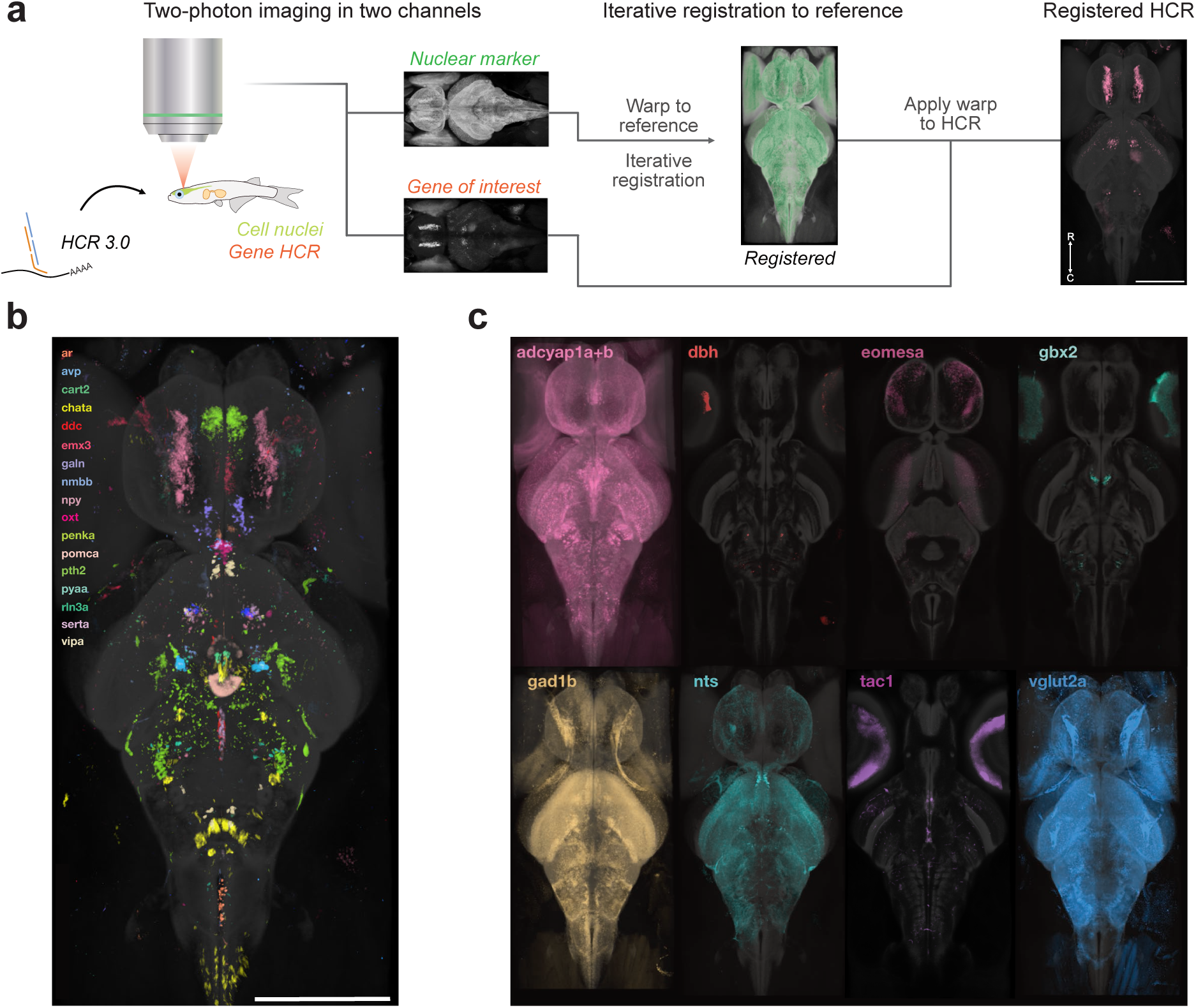
Imaging and registration of neuronal molecular markers. (a) An overview of the staining pipeline: two-photon recordings of whole brains of transgenic fish expressing nuclear GCaMP6s (green) and stained for a neuronal marker using the HCR 3.0 protocol (red). The fish is then registered to the reference brain through the green nuclear channel and the staining is transferred to the reference. R-C, rostral-caudal. (b) A multicolour maximum intensity projection overlay of selected neuronal markers registered to the reference brain with the reference brain in the background. (c) An overview of other stainings included in the atlas, all overlaid on the reference brain. All images are maximum intensity projections apart from *gbx2, dbh, eomesa* and *tac1*, which are shown as slices with staining present due to small labelled populations and high background in the maximum intensity projections. Additional stainings available either in the Supplementary Fig. 4, or in the atlas repository (*isl1a*, *slc6a3, pdyn*). All probes used are listed in the Supplementary Table 1. All scale bars, 500 µm.

**Figure 3.**
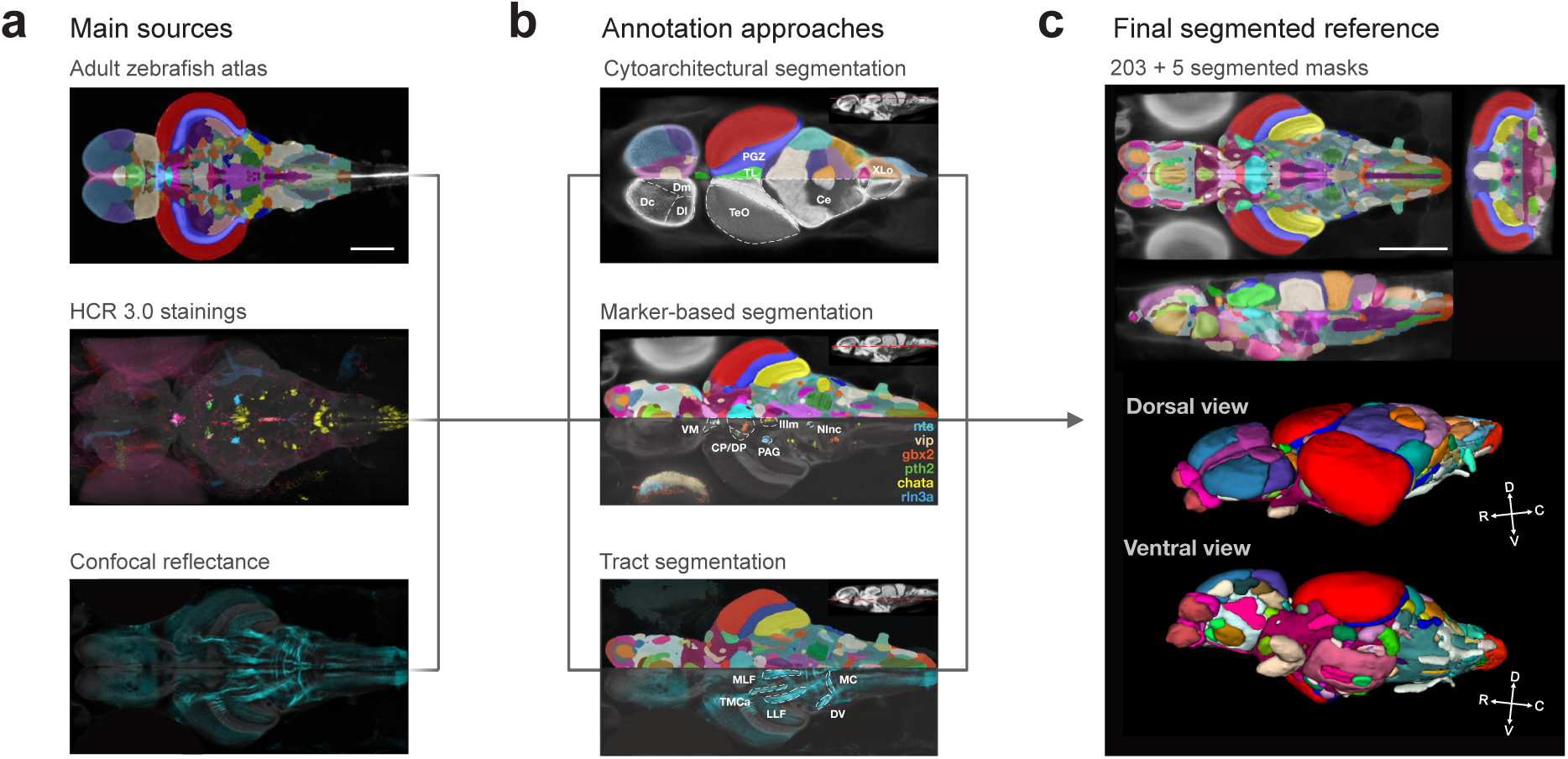
Anatomical segmentation of the *Danionella cerebrum* brain. (a) Resources used as a background for the segmentation: zebrafish annotation from the adult zebrafish brain atlas (AZBA), molecular stainings using HCR 3.0, confocal reflectance imaging of fibre tracts inside the *Danionella brain* (eyes masked). (b) Corresponding annotations that primarily relied on each of the background modalities: cytoarchitectural segmentation based on the nuclear contrast when compared to AZBA; marker-based segmentation of neuronal populations that expressed specific markers; fibre tract segmentation based on negative contrast in the reference and on the confocal reflectance imaging. (c) Final fully segmented volume in orthogonal middle slice views (top) and as a 3D rendered volume using ITK-SNAP in dorsal (middle) and ventral views (bottom). D-V, dorsal-ventral; R-C, rostral-caudal. All brain area abbreviations are listed in the Supplementary Table 2. Scale bars, 500 µm.

### Tissue clearing and reference brain creation

To create a neuroanatomical reference dataset, we used transgenic *Danionella cerebrum* pan-neuronally expressing nuclear-bound GCaMP6s (Hoffmann et al., 2023), exploited here as a bright nuclear fluorescent marker. To visualise the distribution of neuronal nuclei across the entire brain, we fixed and cleared these specimens using a modified passive clarity (PACT) clearing protocol (Supplementary Fig. 1b-c, Yang et al., 2014). We then imaged them using a two-photon microscope at < 1 µm horizontal resolution and with 5 μm axial steps (approximately the size of a cell body). In total we recorded 21 full brain stacks from 11 male and 10 female fish, providing balanced coverage across sexes for template construction (Fig. 1a, Supplementary Fig. 1).

We then used these recordings to create an average reference brain. Given the observed variability in the overall brain shape across individual fish, we chose an iterative non-linear registration approach using the ANTsPy package (Tustison et al., 2021). Our approach consisted of first linearly rotating, translating, and scaling all brains onto each other to create the initial average, followed by iteratively registering all brains to this template using the template building routine with affine and diffeomorphic registration steps with multiple resolutions and smoothing levels (Fig. 1a; see Methods for details). This resulted in a well-resolved average brain that we used as the basis for all subsequent registrations of anatomical, molecular and functional datasets (Fig 1b-c).

### Whole-brain molecular marker labelling

Identification of various neuronal populations and brain areas often relies on the expression patterns of specific neuronal markers. This is especially true in *Danionella cerebrum*, where the small brain and relatively low neuron number even in adulthood (Schulze et al., 2018) reduce the prominence of cytoarchitectural landmarks in nuclear-contrast images. Therefore, molecular marker expression can be a powerful approach to delineate and validate anatomical borders, as many regions in our reference brain appear as small cell groups without clearly distinguishable boundaries (see, e.g., Supplementary Fig. 2). In order to separate and molecularly define these areas, we employed the HCR 3.0 protocol to detect the mRNA distribution of various neuronal markers.

Transcripts for each of the marker genes were stained in a fish pan-neuronally expressing nuclear-localised GCaMP6s to ensure precise registration to the reference (Fig. 2a). We labelled major neuronal cell types: excitatory *vglut2.1+* neurons, inhibitory *gad1b+,* cholinergic *chata+,* as well as neuromodulatory neurons (*ddc*, *serta, dat, dbh).* To distinguish rare neuronal subpopulations we also stained an array of neuropeptides, including *adcyap1, npy, galn, avp, oxt, vipa* and others (Hiraki-Kajiyama et al., 2024). The gene list was iteratively expanded for identification of remaining areas with probes for various transcription factors and other markers (see full list in Supplementary Table 1). We included 29 individual stainings in this initial release of the atlas (Fig. 2b-c), covering major neurotransmitter classes, neuromodulatory systems and neuropeptidergic populations to facilitate the subsequent anatomical segmentation, and established a registration pipeline to add additional labellings in the future.

### Anatomical segmentation

In our delineation of brain areas in *Danionella cerebrum*, we combined whole-mount HCR stainings with comparative teleost neuroanatomy, drawing primarily on adult zebrafish resources (Anneser et al., 2024; Hiraki-Kajiyama et al., 2024; Mueller, 2012; Porter & Mueller, 2020). Most detailed teleost datasets, however, are based on densely sliced transverse sections, whereas our whole-mount preparations were imaged in horizontal planes, complicating a direct one-to-one comparison of anatomical borders with the transverse slices in the literature. To segment anatomical regions in three dimensions despite these differences in orientation, we thus used the adult zebrafish brain atlas (AZBA, Fig. 3a, top) as a guiding reference (Kenney et al., 2021), comparing Danionella cytoarchitecture and molecular marker patterns to AZBA. This resource transfers and updates the anatomical annotations from the zebrafish neuroanatomical atlas (Wullimann et al., 1996), and we therefore additionally adopted its approach to brain area hierarchy and terminology as the starting point for our *Danionella cerebrum* segmentation.

We initially annotated the most prominent structures – parts of the optic tectum and cerebellum, the habenula, large cranial nerve lobes and their corresponding cranial nerves – based on their clear anatomical borders and similarity to zebrafish (Fig. 3b, top). These structures served as anchor regions. In the next step, we iteratively expanded the segmentation around these anchors, identifying and confirming further regions with the available stainings (Fig. 3b, middle panel). Finally, we refined and adjusted the borders of all annotated regions using the stainings until we achieved a mutually consistent segmentation across the different orthogonal planes (Fig. 3c).

In the hindbrain, most of the prominent nuclei were annotated visually; *chata* and *isl1a* stainings allowed us to find the efferent octavolateral (OENr and OENc) and trigeminal motor nuclei (Vmv) (Baeuml et al., 2019; Brantley & Bass, 1988; Lozano et al., 2023) (Supplementary Fig. 2a). We used the *dbh* to find the locus coeruleus (LC), *serta* for superior (SR) and inferior raphe (IR), in addition to *adcyap1* expression in r the isthmic nucleus (NI) and *ar* in the IXth cranial nerve motor nucleus (IXm) that we identified based on cytoarchitecture (Hu et al., 2002; Pérez et al., 1996). The medullary nucleus incertus (NInc) belonging to the caudal central grey (GC) was identified based on cytoarchitecture and confirmed with neuropeptides *rln3a* and *nmbb* (Spikol et al., 2023) (Supplementary Fig. 2b). Remaining nuclei in the hindbrain and spinal cord were annotated by comparing them to the zebrafish hindbrain in AZBA and the zebrafish neuroanatomical atlas (Wullimann et al., 1996).

Midbrain optic tectum (TeO and PGZ), as well as tegmental torus longitudinalis (TL), torus semicircularis (TSc), medial pretoral nucleus (MPN), and interpeduncular nucleus (NIn) were identified based on their cytoarchitecture and served as anchors for the rest of the annotation. Like in zebrafish, strong *chata* staining identified the oculomotor nucleus (IIIm) separating nucleus of the medial longitudinal fascicle (NMLF) from the MLF (Supplementary Fig. 2a). In addition, we confirmed the location of the NMLF through a spinal backfill injection with a tracer dye (Supplementary Fig. 3a) (Akbari et al., 2024; Kimmel et al., 1982). We verified the rostral extent of the midbrain central grey by finding the *rln3a-*expressing subpopulation (that we labelled as the genetically defined rostral PAG (rPAG), see (Spikol et al., 2023)) adjacent to the *nmbb-*expressing Edinger-Westphal nucleus (EW), tentatively annotated based on *nmbb* expression shown in mammals (Burnell et al., 2008; Priest et al., 2023) (Supplementary Fig. 2b). We separately annotated the auditory medial pretoral nucleus (MPN) rostral to the torus semicircularis (TS) (Northcutt, 2006; Yamamoto & Ito, 2005) as part of the ascending auditory pathway.

In the diencephalon, we identified the central and dorsal posterior thalamic nuclei with the expression of the transcription factor *gbx2* (Mallika et al., 2015; Wullimann et al., 2024) and the ventromedial thalamus (VM) through high expression of the neuropeptides *nts* and *vipa* (Bello et al., 1994; Hiraki-Kajiyama et al., 2024, Supplementary Fig. 2c, top), with the rest of the thalamic divisions identified anatomically with reference to AZBA. Anterior (PPa) and posterior (PPp) preoptic areas were identified anatomically and confirmed through expression of *galn* in the anterior part (Amano et al., 2009), with the magnocellular (PM) portion visible through the expression of the neuropeptides *avp* and *oxt,* which play important role in the regulation of social behaviours (Eaton et al., 2008; Goodson, 2008, Supplementary Fig. 2c, bottom). For the annotation of the posterior tuberculum and hypothalamus, we relied on anatomical similarity with the AZBA and confirmed specific borders through comparisons with the zebrafish neuropeptide atlas (Hiraki-Kajiyama et al., 2024), yielding a diencephalic segmentation closely aligned with zebrafish nomenclature. In addition, we have identified the pituitary (Pit) through the expression of *pomca* and saccus vasculosus (SV) based on its cytoarchitecture ventral to the hypothalamus (Nakane et al., 2013; To et al., 2007).

Telencephalon annotation presented the largest challenge given the sparse cellular density in the pallium without distinct borders, and the compact size of the subpallium. We used markers identified in zebrafish (Ganz et al., 2015) to support the identification of the pallial divisions (*eomesa* for Dc and Dl, *emx3* for Dm and *emx1* for Dp, Supplementary Fig. 2d, top). Given the relatively low expression of these genes detected in our hybridisations, we additionally injected a dextran tracer dye into the olfactory bulb (Supplementary Fig. 3b) to confirm the location of the dorsoposterior pallium (Dp), with strongly labelled olfactory projections identified ventral to the dorsolateral Dl and dorsomedial Dm (Kermen et al., 2013). Additionally, this allowed us to visualise the medial and lateral olfactory tract (MOT). In the ventral telencephalon, we identified the subpallium and separated the dorsal and central parts (Vd/Vc) from the ventral (Vv) through the expression of *penka* and *tac1* (Hegarty et al., 2024; Hiraki-Kajiyama et al., 2024; Tanimoto et al., 2024b; Tibi et al., 2023, Supplementary Fig. 2d, bottom). The rest of the nuclei present in AZBA were annotated based on the cytoarchitectural similarity to zebrafish, as well as adjacent nuclei that were identified through stainings and tracing.

Finally, we identified the nerves and tracts present in AZBA within the *Danionella* brain. Because the contrast in our reference brain is nuclear, template averaging rendered large tracts as low-intensity regions, allowing easy visual identification through negative contrast. For the finer borders and smaller tracts, we additionally acquired a 3D image of a live fish under two-photon microscope with a confocal reflectance channel, which allowed us to directly visualise large myelinated tracts that backreflected the excitation light (Fig 3b, bottom, Pimentel-Domínguez et al., 2025). Together with the specific stainings (such as *npy* associated with the MOT (Gaikwad et al., 2004) and the olfactory bulb injection (anterior commissure, AC), we placed all of the fibre tracts annotated in zebrafish in our atlas. As a result, we were able to generate a mutually exclusive and collectively exhaustive (MECE) annotation set for the entire *Danionella cerebrum* brain (Fig. 3c). All segmented areas, together with the marker stainings and references used to support their identification, are listed in Supplementary Table 2.

### Cross-registration of functional and anatomical imaging

The small size of *Danionella cerebrum* combined with the lack of dorsal skull allows for large-field imaging of neuronal activity using volumetric imaging techniques (Hoffmann et al., 2023). To supplement the molecular data in this brain atlas, we exposed multiple animals to visual stimuli while recording their whole-brain calcium activity using oblique plane microscopy (OPM), and reanalysed OPM data of sound-evoked brain activity from (Henninger et al., 2026) (Fig. 4a, see Methods for details). To bridge the functional recordings to the anatomical reference, we used the OPM-recorded fish to create functional brain templates as a bridge between functional maps and the reference brain. We then transferred the coordinates of active cells to the reference coordinates using the same ANTs-based registration pipeline as for the anatomical data (see Methods, Fig. 4b,d). We then quantified the density of stimulus-responsive cells in each annotated region (Fig. 4c,e).

**Figure 4.**
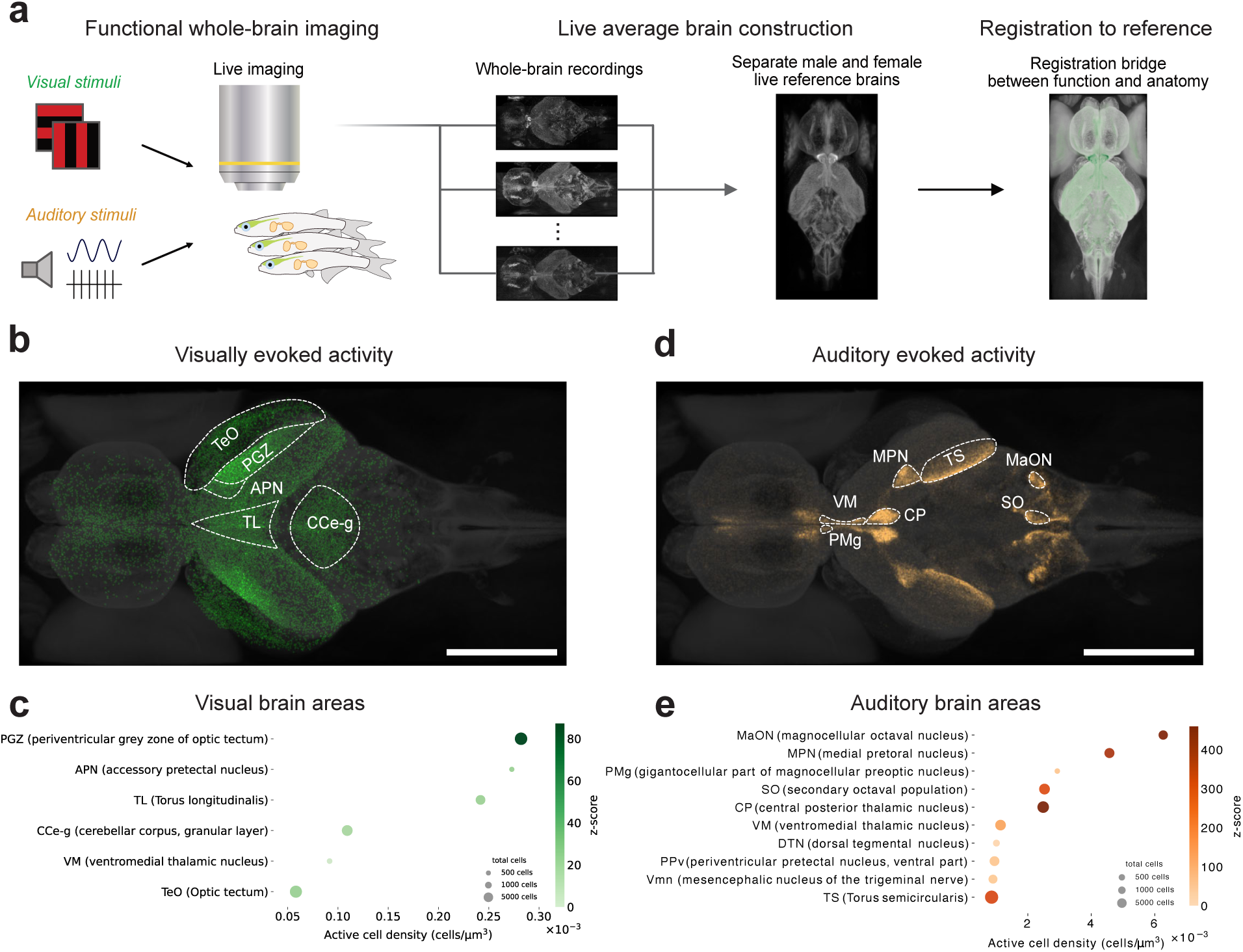
Registration of whole-brain stimulus-evoked functional activity recordings to the reference brain. (a) Overview of the experimental pipeline: first the fish brains were recorded under the microscope during either visual or auditory stimulation. Recorded brain activity was used to create intermediate average template brains in the functional recording space that was then registered and transferred to the reference brain coordinates. (b) Maximum intensity projection of the density of visual stimulus-responsive cells in the brain of Danionella cerebrum overlaid on the reference template. Number of fish, 6. (c) Brain areas with significantly overrepresented visual stimulus-responsive cells sorted by responsive cell density. (d) Maximum intensity projection of the density of auditory stimulus-responsive cells in the brain of Danionella cerebrum overlaid on the reference template. Number of fish, 44. (e) Top 10 brain areas with significantly overrepresented visual stimulus-responsive cells sorted by responsive cell density See Supplementary Fig. 4 for a longer list. (c) and (e) significance: p < 0.05 permutation testing with per-fish stratification (see Methods). Z-score indicates the deviation of the observed cell distribution from the permutation-based null distribution. Scale bars, 500 µm.

As expected, visual stimuli were mostly correlated with neuronal activity in parts of the optic tectum (TeO, PGZ), including the accessory pretectal nucleus (APN) that is associated with prey capture (Antinucci et al., 2019; Semmelhack et al., 2014) and the longitudinal torus (TL) involved in the binocular integration (Tesmer et al., 2022) (Fig. 4b,e)). In addition, visual responses were observed across the granular layer of the cerebellum that has been shown to integrate sensorimotor information including vision (Knogler et al., 2017). Responses to the auditory stimuli were prominent across the entire auditory network (Fig. 4d,e; Supplementary Fig. 4), including in the octavolateral system (descending octaval nucleus (DON), anterior octaval nucleus / magnocellular nucleus (AO/MaON) and the secondary octaval population (SO), (Edds-Walton et al., 2010; Maruska & Tricas, 2009; McCormick & Braford, 2009)), the auditory midbrain (medial pretoral nucleus MPN and torus semicircularis TS, (Ma & Fay, 2002)), the auditory thalamus (central posterior thalamic nucleus CP, (Kirsch et al., 2002)), and the diencephalon (posterior tuberculum (TPp) and preoptic area (PMg), (Forlano et al., 2017; Perelmuter et al., 2019)).

In summary, these functional data confirmed the anatomical identification of the nuclei known to be involved in visual and auditory sensory processing in teleosts and validated our segmentation in functionally defined circuits. We deposited these registered maps as part of the atlas to guide future experiments involving these modalities and to provide a first functional layer for the *Danionella cerebrum* atlas.

### Sexual dimorphism in the brain of *Danionella cerebrum*

Whereas adult zebrafish atlases such as AZBA reported no substantial sex-dependent differences in overall brain anatomy (Kenney et al., 2021), pronounced sexual dimorphism in brain morphology is well documented across vertebrates. Examples from teleosts include differences in overall brain size (sticklebacks), dorsolateral pallium organisation (blenniids), and pituitary structure (medaka) (Costa et al., 2011; Kotrschal, Räsänen, et al., 2012; Royan et al., 2021). *Danionella cerebrum* displays pronounced sexual dimorphism in body morphology and behaviour (Britz et al., 2021; Groneberg et al., 2024; Henninger et al., 2026; Penalva-Tena et al., 2025). We therefore asked whether male and female Danionella also differ in brain size and regional anatomy, and quantified these differences.

To examine sex-dependent differences in brain size and regional anatomy, we registered individual 11 male and 10 female brain volumes used in the creation of the reference brain to the reference brain itself. Global linear scaling analysis revealed that female fish brains were significantly larger than male brains (mean±std brain volumes; female, 802.16±103.18 nL, male, 649.94±81.19 nL; volume ratio 0.81, p-value, 0.003, Welch t-test). Afterwards, we quantified the local warping during the registration step (Fig. 5a). Average reference brains generated separately for both sexes confirmed gross morphological differences most evident in the telencephalon and cerebellum (Fig. 5b). When the relative volume size for each region was quantified (Fig. 5c), female brains showed larger cerebellar structures: 30% larger cerebellar crest CC, 25 % larger molecular layer of the lateral valvula (Val-m), and 21% larger caudal lobe (LCa), as well as separate divisions of the octavolateral system: 21% larger medial (MON) and 19% larger caudal (CON) octaval nuclei. Male brains had 71% larger lateral dorsal pallium (Dl), as well as 66% larger lateral preglomerular nucleus (PGl) and 50% larger facial lobe (VIILo). These differences are presented on a per-voxel comparison effect size map of differences seen in Fig. 5d and provide an anatomical basis for future studies of sexually dimorphism in *Danionella cerebrum*.

**Figure 5.**
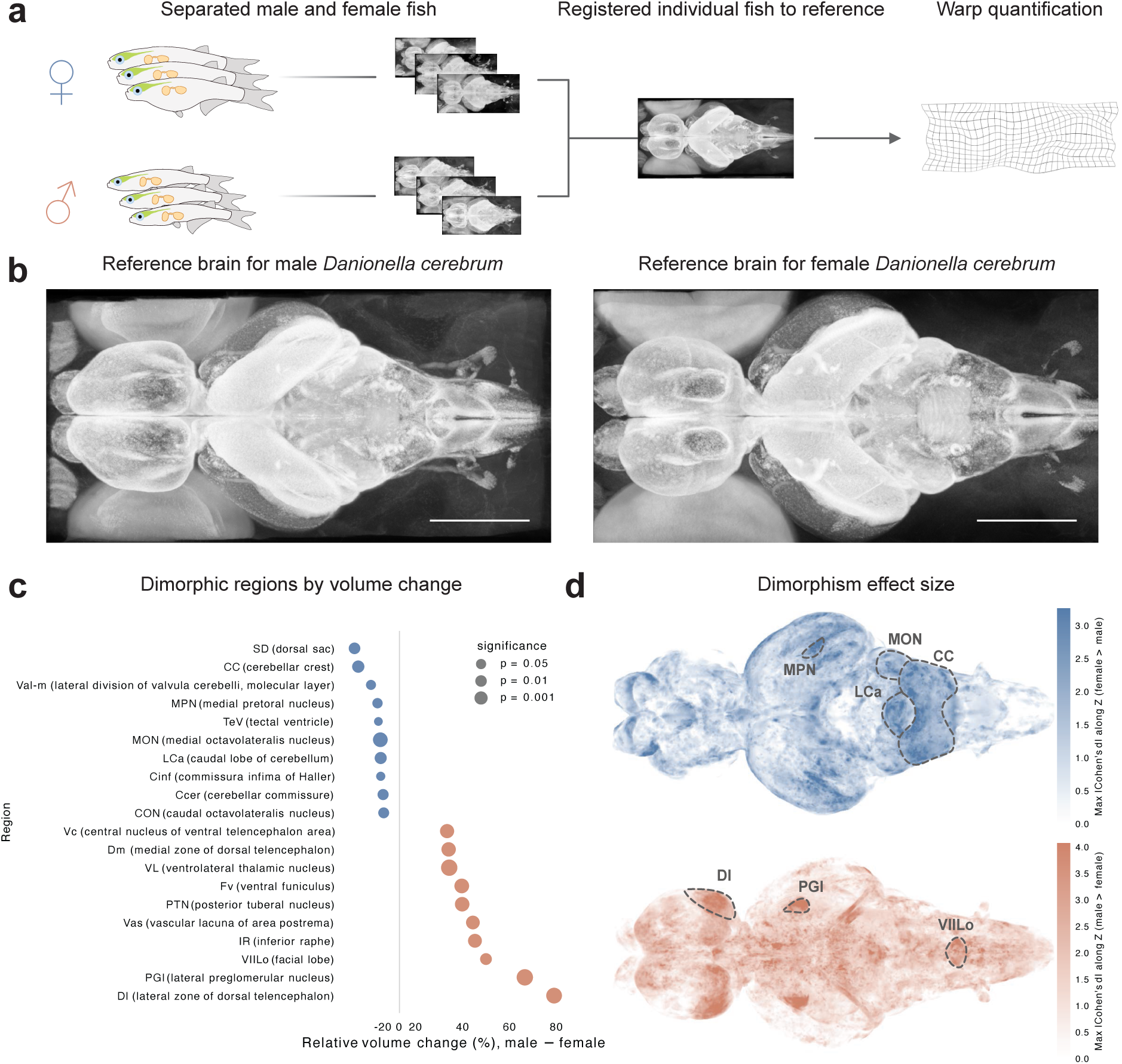
Sexually dimorphic brain morphology in *D. cerebrum*. (a) Male and female fish brains used for template constructions are separately registered to the reference brain to generate expansion and contraction maps using Jacobian determinants of the warp fields. (b) Reference brains created from male (left) and female (right) brains only to showcase the large-scale differences in brain morphology. (c) Quantification of the top 10 most dimorphic brain regions by relative volume difference in female and male fish each. Blue indicates areas larger in females and red indicates areas larger in males. A two-sample Welch t-test per area. (d) Effect sizes of per-voxel differences across the entire brain, represented as maximum intensity projections of absolute Cohen’s d values for the log-Jacobian determinants of the warp fields. Red indicates areas larger in males, blue indicates areas larger in females.

### Atlas structure, access and workflow

To make the *Danionella cerebrum* atlas a lasting community resource capable of incorporating new data types and updated annotations, we organised it as a versioned GIN repository (https://gin.g-node.org/danionella/dc_atlas; Herz et al., 2008), a git-based neuroscience data management platform. The repository combines reference atlas with registered datasets and dataset-level metadata in a unified structure (see Data Availability statement). Structural, molecular, and functional datasets are stored in an expandable format and linked through standardised human-readable metadata. A registration tutorial and contribution guideline provide clear instructions for adding new datasets and maintaining compatibility across versions.

The resources deposited with the atlas are viewable through standard software such as ITK-SNAP (Yushkevich et al., 2016). In addition, we created and hosted a web viewer (https://atlas.danionella.org/cerebrum) for rapid exploration of the datasets described here and those added in the future. Together, this structure provides a workflow for accessing, visualising, and expanding the atlas.

## Discussion

We present a neuroanatomical resource for *Danionella cerebrum* to facilitate its application as a model organism for neuroscience. We optimised tissue clearing, RNA amplification, and imaging protocols to achieve whole-brain high-resolution anatomical imaging, neuronal marker transcript detection, and functional activity mapping. We created an average reference brain and showed its applicability as a common coordinate system for mapping experimental data across imaging modalities. 29 individual neuronal marker stainings combined with our reference brain based on nuclear labelling-derived anatomical contrast allowed us to segment and annotate 203 anatomical areas that correspond to the updated zebrafish resources.

Our whole-mount staining protocol was capable of identifying main populations of interest for all included neuronal markers; however, markers that are more challenging to visualise, including transcription factors and receptors such as *gbx2* and *ar*, as well as subpopulations that express selected markers weakly, such as *pth2* in PGZ or *chata, emx1 and emx3* in the telencephalon, were not clearly identifiable across their entire previously published expression patterns even with repeated stainings. Future development of the atlas, including further optimised and averaged molecular labellings across multiple specimens and improved and more sensitive staining protocols such as HCR Gold, is likely to expand the set of identified populations and refine borders between neighbouring nuclei. Integration with presently available spatial transcriptomics datasets (Atabay et al., 2026), as well as multiplexed whole-mount stainings combined with these more sensitive protocols could reveal additional detail to the borders of areas that are identified based on the expression of multiple markers, for example, *shh-*expressing dopaminergic cells at the border area between the ventral posterior tuberculum and hypothalamus (Wullimann & Umeasalugo, 2020). Furthermore, the addition of further neurodevelopmental markers (e.g. patterning genes and transcription factors with neuromeric specificity) will allow us to incorporate more recent neuroanatomical developments that reframe diencephalic nuclei within the genoarchitectonic, neuromeric model (Wullimann, 2022). Thus, the anatomical segmentation and the underlying stainings presented in this study are a first step towards a living anatomical resource that could be continually updated with new information in the future, which we facilitate by structuring it as a versioned metadata-rich resource.

Our study reveals that sexual dimorphism in *Danionella cerebrum* extends beyond the body and male-specific drumming muscle morphology (Britz et al., 2021; Groneberg et al., 2024) to the brain itself, with pronounced differences across telencephalic, cerebellar and other hindbrain nuclei. These findings raise the questions of how the neuroanatomical differences relate to sex-specific behaviour and physiology (Bass & Baker, 1990; Feng & Bass, 2016). Some of the apparent dimorphism in the dorsal pallium might partly reflect variation in the anatomical position of the brain relative to the eyes; however, dorsolateral pallium has previously been described as sexually dimorphic in blenniid fish (Costa et al., 2011). Given its role in spatial learning and navigation in teleosts (Fotowat et al., 2019), such dimorphism may reflect sex-specific navigational demands, consistent with reports of a relatively larger cerebellum in female guppies (Kotrschal, Rogell, et al., 2012). Differences in hindbrain sensory-motor structures may likewise relate to sex-specific behavioural states: the facial lobe shows seasonal variation in reptiles (O’Bryant & Wade, 1999). Aromatase expression and catecholaminergic innervation of octavolateral nuclei varies seasonally in female midshipman fish (Forlano et al., 2015; Forlano & Bass, 2005), suggesting that reproductive context can reshape circuits related to sensory processing and social communication. Studying how these and other features are shaped throughout development could shed light on how steroid hormones regulate neurodevelopment and result in sexually dimorphic brain functions and behaviours.

In summary, the *Danionella cerebrum* brain atlas provides a framework for integrating anatomical, molecular, and functional data in an optically accessible adult vertebrate brain. Together with the integration of datasets from across the *Danionella* community, it will support future work on neuroanatomy, physiology, gene function, and sexual dimorphism in this model organism.

## Methods

### Animal husbandry and transgenic lines

All animal experiments and husbandry conformed to Berlin state, German federal and European Union animal welfare regulations. *Danionella cerebrum* were housed as previously described (Groneberg et al., 2024) in water circulated aquaria (Tecniplast) supplied with artificial fish water (Instant Ocean at a conductivity of 350 μS/cm, pH 7.3 adjusted using NaHCO_3_) at 26-27 °C and with a 14:10 h light:dark cycle, with lights on from 08:00 until 22:00 h preceded by a 30 min dimming period. We used adult (∼6-12 months old) *Danionella cerebrum* expressing nuclear-bound GCaMP for the construction of the template, functional imaging, and as the reference channel in molecular stainings. For all reference brains and majority of the stainings, a tyrosinase knock-out line crossed with a pan-neuronal nuclear-bound GCaMP6s reporter line (either *tyr^-/-^* ⨉ elavl3:GCaMP6s or *tyr^-/-^* ⨉ snap25a:GCaMP8m with similar but denser expression pattern) was used to remove melanin pigmentation on top of the head. For functional imaging experiments with visual stimulation and for some of the stainings (*vglut2.1, gad1b*), a MITF knockout line (*mitfa^-/-^)* crossed with the GCaMP reporter was alternatively used to prevent depigmentation of the eye and resulting high eye autofluorescence, respectively (Johnson et al., 2011).

### Sample preparation, fixation, and tissue clearing

For anatomical imaging, the fish were euthanised by submersion in ice water, placed in ice-cold PBS with 0.5 M EDTA and decapitated using a fine scalpel. Afterwards they were fixed in a 4% formaldehyde (Pierce) solution in PBS for 24 hours at 4℃. The next day the samples were washed 2 ⨉ 30 minutes in PBST (PBS with 0.1% Tween-20 added, Sigma). Then the dorsal skin covering the brain was gently pierced and partially removed using fine 5SF forceps (Fine Science Tools) to improve the diffusion of the clearing solution and staining reagents into the brain. Afterwards, the samples were placed into 1.5 mL of clearing and permeabilisation solution (1X PBS, 1% Triton X-100, 0.2% SDS, 20 mM sodium sulfite) at 25℃ for 48 hours with gentle shaking. For larger female fish, the temperature was adjusted to 37℃. After the clearing, the samples were washed for at least 4 hours in PBST. Fish used for molecular marker staining were additionally stained as described below, while transgenic fish used for the reference creation were immediately placed into a refractive index matching solution (RIMS, EasyIndex custom RI 1.465, LifeCanvas Technologies).

### HCR 3.0 staining of neuronal markers

For all HCR 3.0 stainings, B1-Alexa 594 HCR amplifiers were used with the corresponding probe sequences. Permeabilised samples were washed in 1X PBS for 4 hours and stained using a modified HCR protocol for whole-mount samples (Lovett-Barron et al., 2020). After 2 ⨉ 20 minute 2X SSCT (2X SSC, 0.1% Tween-20) washes, the hybridisation probes were added to each fish at 1:250 dilution in fresh hybridisation buffer (2X SSC with 10% v/v formamide (Sigma) and 10% v/v dextran sulfate (50% solution, Sigma)). The hybridisation was conducted at 37℃ for 18-22 hours. Afterwards, the samples were washed 3⨉ in a probe wash solution (2X SSCT with 30% formamide) at 37℃ and 2⨉ in 2X SSCT at 24℃, 20 minutes each. Afterwards, they were incubated in the amplification buffer (5X SSCT with 10% v/v dextran sulfate). Meanwhile, the Alexa Fluor 594-bound fluorescent amplifiers were denatured at 95℃ for 1 minute in a PCR machine and then immediately snap cooled on ice. They were then allowed to return to room temperature for 20 minutes in a dark drawer and subsequently added to the samples in a fresh amplification buffer (4 µL of each hairpin added to 50 µL of amplification buffer per fish). The samples were left at room temperature overnight. The next day they were washed 3 ⨉ 30 minutes in 5X SSCT and finally optically cleared in ∼300 µL of RIMS for 24 hours.

### Two-photon anatomical and neuronal marker imaging

Samples were placed in custom Sylgard holders, suspended in RIMS and covered with a coverslip. They were then imaged using a water-immersion Nikon 16X NA 0.8 objective on a custom-built tunable-laser (Chameleon Vision-S, Coherent) two-photon microscope equipped with green (525/50, Edmund Optics) and red (607/70, AHF analysentechnik AG) channels. Image acquisition was controlled via ScanImage 5.6.1 and MATLAB 2019a. For the anatomical GCaMP recordings used for the reference brain construction, we used 920 nm excitation wavelength. To record both reference GCaMP signal and the HCR staining signal from Alexa Fluor 594, we used 820 nm excitation wavelength, which was a trade-off between the signal in both channels and the sample autofluorescence. Samples were imaged in tiles with a 617 nm lateral pixel pitch and 5 μm axial steps, covering the entire brain over a ∼2.5 × 1 × 0.6 mm^3^ volume.

### Brain reference template creation and registration

Acquired images were stitched using the Fiji stitching plugin (Preibisch et al., 2009; Schindelin et al., 2012). For molecular stainings that showed high amount of blood vessel autofluorescence, we used the fact that HCR amplicons were present in the cytosol and GCaMP was bound to the nucleus to normalise and subtract the GCaMP channel from the molecular staining channel, thus mostly removing the autofluorescence present in both channels. The contrast in the GCaMP reference channel was then enhanced using a CLAHE algorithm implemented in Fiji. All further processing and registration has been done using ANTsPy in Python (Tustison et al., 2021). All stacks were downsampled to a ∼2.5 ⨉ 2.5 ⨉ 5 µm^3^ voxel size (X ⨉ Y ⨉ Z) to speed up the registration.

To generate the reference brain template, the 21 individual brains stacks were initially pre-aligned and pre-scaled to a set average size with the AntsPy ‘Similarity’ registration routine. Their arithmetic average served as an initial template. It was then iteratively refined over 4 iterations using the AntsPy build_template routine with ‘SyNRA’ (rigid, affine, then symmetric normalisation) registration, 200 iteration per stage, with (4, 3, 2, 1) smoothing and (12, 8, 4, 1) shrinking stages. The same procedure was applied when constructing the sex-specific templates using only female or male fish brains, as well as constructing the templates for the functional imaging modality.

All other anatomical and molecular modalities were registered to the reference brain using the reference GCaMP channel. Initially, the fish were preregistered using the ‘TRSAA’ (translation, rotation, scaling, 2x affine) routine to align them to the reference brain as closely as possible, followed by more stringent ‘SyNCC’ registration routine (affine and symmetric normalisation with cross-correlation as a metric of volume similarity) with default parameters as of ANTsPy version 0.3.8.

For a subset of brains, the default ‘SyNCC’ registration parameters produced strong overfitting and misalignments in the telencephalon (e.g., *cart2, gad1b, eomesa, isl1a, pdyn,* and *vglut2a* stainings). For these cases, we instead applied a two-step ‘SyNOnly’ registration strategy. First, the pre-aligned brains were conservatively warped with four multi-resolution stages with maximum iterations of (250, 200, 150, 80), SyN metric sampling 8, gradient step 0.10, flow sigma 3 and total sigma 1.5. Afterwards, the registrations were refined using a more aggressive SyN pass with (300, 250, 200, 150) iterations, SyN metric sampling 6, gradient step 0.12, flow sigma 3 and total sigma 0.5. To reduce spurious deformations outside of the brain, we additionally created masks in both moving and fixed spaces and restricted the optimisation to the brain volume.

### Anatomical segmentation

All brain areas were segmented manually on the reference brain using ITK-SNAP (Yushkevich et al., 2016) with the stainings and confocal reflectance channel overlays to inform the segmentation. We used the AZBA zebrafish neuroanatomical segmentation for the terminology and as our guide for the clearly visible areas and nuclei. For sparser and small nuclei that weren’t clearly visible in *Danionella* we used available stainings first and then annotated the rest by combination of their vicinity to the known zebrafish nuclei and anatomical similarity. The fibre tracts were annotated using the intensity drops along the fibres in the nuclear-contrast reference brain and the confocal reflectance channel as the contrast for larger tracts. Finally, remaining unclear borders in any given plane were conservatively annotated by the principle of exclusion. Afterwards, the space between neuroanatomical areas has been filled with the “Remaining” brain area masks approximately divided across major brain divisions to create a whole-brain mutually exclusive and collectively exhausting (MECE) mask. All labels were smoothened after the annotation and later individually after each subsequent update using ITK-SNAP recursive gaussian label smoothing feature with a 2⨉2⨉2 voxel standard deviation.

### Functional imaging and registration to the reference

Before the functional imaging, fish were immobilised as previously described (Hoffmann et al., 2023). They were anaesthetised in 120 mg/L MS-222 and then transferred to a custom-made agar mould, where they were perfused with oxygenated water supplied by a peristaltic pump through a pipette inserted into the mouth and then embedded with a few droplets of a low-melting point agarose solution (Fisher Scientific). They were then placed under the objective of an oblique plane microscope (OPM; Henninger et al., 2026; Hoffmann et al., 2023). All specimens were imaged at 1 Hz volumetric rate.

To record visually evoked responses, we used six adult male *Tg(mitfa^-/-^⨉ elavl3:H2B-GCaMP6s)* fish. Each fish was placed at the center of a cylindrical tank (outer diameter: 110 mm; inner diameter: 100 mm; height: 80 mm). A flexible 6 inch AMOLED display (2880 × 1440 px; 35 Hz refresh rate; Wisecoco) was wrapped around the tank, covering approximately 120° of the visual field in front of the fish. The stimulation protocol began with 5 min of a black screen baseline, followed by 30 presentations of static visual stimuli (10 s each), separated by 20 s black-screen interstimulus intervals. Stimuli consisted of vertical and horizontal black-and-red bar patterns with a spatial period of 10 mm, as well as random black-and-red pixel patterns. Each stimulus type was presented 10 times in pseudorandom order.

For acoustically-evoked activity, we reanalysed the datasets for 44 adult fish from (Henninger et al., 2026; see the original publication for detailed methods). In brief, two different types of sound were presented: sequences of short pulses (< 1 ms) that mimic the vocalizations of *Danionella cerebrum* at repetition rates between 4 and 1000 Hz and lasted between 9 ms and 4 s, and pure tones with frequencies between 150 Hz and 2500 Hz at a fixed duration of 2 s. All sounds were presented 5 to 18 times in pseudorandom order. Each trial epoch lasted 15 s to allow the sound-evoked signal to recede before the next sound was presented.

We registered each recording to common, sex-specific reference templates. To generate these templates, we first calculated an average volume for each recording based on the 100 best correlating (r=0.920) volumes. These average brains were then used to create male and female reference OPM templates. Due to the larger size of female fish this step resulted in partial clipping on the sides of the optic tectum. Next, the OPM templates were registered to the corresponding male and female anatomical templates to bridge them to the reference brain.

Individual recordings were then analysed using the OPM processing pipeline established in our lab to correct for motion artifacts and extract cells in the original fish space (Henninger et al., 2026; Hoffmann et al., 2023). Averages of these recordings were then registered to the OPM templates using the ‘SyNCC’ routine as previously described with an added *totalFieldVarianceInVoxelSpace=1* warp penalty that prevented cells from smearing during registration (*--transform SyN[0.15,3,1]*) and then transferred through the previously registered OPM-to-reference affine and warp maps to the reference brain. These transformations were then used to identify coordinates of functionally active cells in the reference coordinate system.

### Confocal imaging of neuronal tracts

For the imaging of fibre tracts, a live transgenic *Danionella cerebrum* was embedded as described above and a whole-brain stack was acquired in two channels: the green channel for the GCaMP expression and the additional confocal reflectance channel that used the back-scattered light through a confocal pinhole onto a silicon photodiode (ThorLabs). This allowed us to image larger myelinated tracts due to their high fat content (Figure 3a, bottom). This confocal reflectance channel was registered to the reference brain through the GCaMP channel as described above.

### Tracer injections

The GCaMP fish were anaesthetised as described above and placed on an agarose mould under a stereoscope (Olympus). Fixable Alexa 594-dextran (MW 10k, Thermofisher) was injected either into the olfactory bulb unilaterally or into the spinal cord (Supplementary Fig. 3a,b) through a 5 μm injection pipette (Clunbury Scientific) using a PicoNozzle with the PicoPump vacuum pressure injector (World Precision Instruments) and the fish were allowed to recover. The next day, they were sacrificed and fixed. After fixation and rinsing, they were immediately placed into the EasyIndex RIMS solution for 1-2 days and imaged.

### Brain area overrepresentation analysis

To assess which anatomical regions were enriched for functionally responsive cells (Fig. 4c,d), we first registered all detected cells from each fish to the common reference template and assigned an anatomical label to each cell (i.e., a brain region where this cell is located). For every brain region *i* we counted the total number of detected cells *n_i_* and the number of “responsive” cells *k_i_* (stimulus-locked or high-effect cells). We then identified brain areas that contained significantly more responsive cells than expected from random distribution of such cells across the brain using a permutation test that preserved fish-to-fish variability as follows. For each permutation we shuffled the responsive cell labels within each fish separately, keeping the total number of these cells *K_u_* and the per-region cell counts within that fish fixed. For each permutation we recomputed the number of responsive cells per region, yielding a null distribution for *k_i_* that respects the observed sampling of cells across animals and areas. We performed 10^6^ such permutations and estimated the one-sided enrichment p-values as the upper-tail probability *Pr*(*K*_*i*_^*perm*^ *k*_*i*_^*obs*^) with Phipson–Smyth bias correction (Phipson & Smyth, 2010). The resulting p-values were adjusted for multiple testing across regions using the Benjamini–Hochberg procedure. To reduce sensitivity to minor registration errors and extremely small structures, we excluded all tract and ventricular labels and discarded regions with fewer than 50 detected cells in total after statistical testing.

### Sexual dimorphism analysis

For the male–female comparisons, we first registered each individual brain to the reference brain template as previously described using an initial linear (affine) stage to remove the linear scaling between different brains followed by a non-linear ‘SyNCC’ warp. We then computed log-Jacobian determinant images from the resulting deformation fields, which quantify the local logarithmic volume change at each voxel relative to the template. To visualise the spatial distribution of sexually dimorphic brain areas (Fig. 5d), we computed a voxelwise standardised effect size (Cohen’s *d*) of the log-Jacobian values between male and female fish and displayed these as maximum intensity projection maps. To identify the most dimorphic brain areas (Fig. 5c), we summarised the log-Jacobian values at the level of annotated regions. For each subject and each brain area, we computed the mean log-Jacobian across all voxels belonging to that label and compared these regional means between males and females using a two-sample Welch t-test per area. Resulting p-values were corrected for multiple comparisons across regions using the Benjamini–Hochberg procedure. For interpretability, we converted the regional differences in mean log-Jacobian back into linear percentage volume changes, such that a value of +100 % corresponds to a region that is, on average, twice as large in males as in females, and negative values indicate female-biased volume increases.

## Data availability

The atlas resources (brain volumes used in the reference creation, as well as the molecular, functional, and structural datasets) have been deposited as a *Danionella cerebrum* atlas repository on G-Node (https://gin.g-node.org/danionella/dc_atlas). Supplementary tables, analysis code, and the associated data have been separately deposited on G-Node (https://gin.g-node.org/danionella/Kadobianskyi_et_al_2026).

## Acknowledgements

We thank J. Perelmuter, A. Bass, A. Douglass, and M. Lovett-Barron for helpful discussions and for feedback on the manuscript. We also thank our fish facility team for excellent fish care and experimental support. We acknowledge support by the European Research Council (ERC2021-CoG-101043615), the Einstein Foundation (EPP-2017-413), the German Research Foundation (DFG, projects EXC-2049-390688087, 432195732 and 532521431) and the Alfried Krupp von Bohlen und Halbach Foundation.

## Supplementary figures

**Figure S1.**
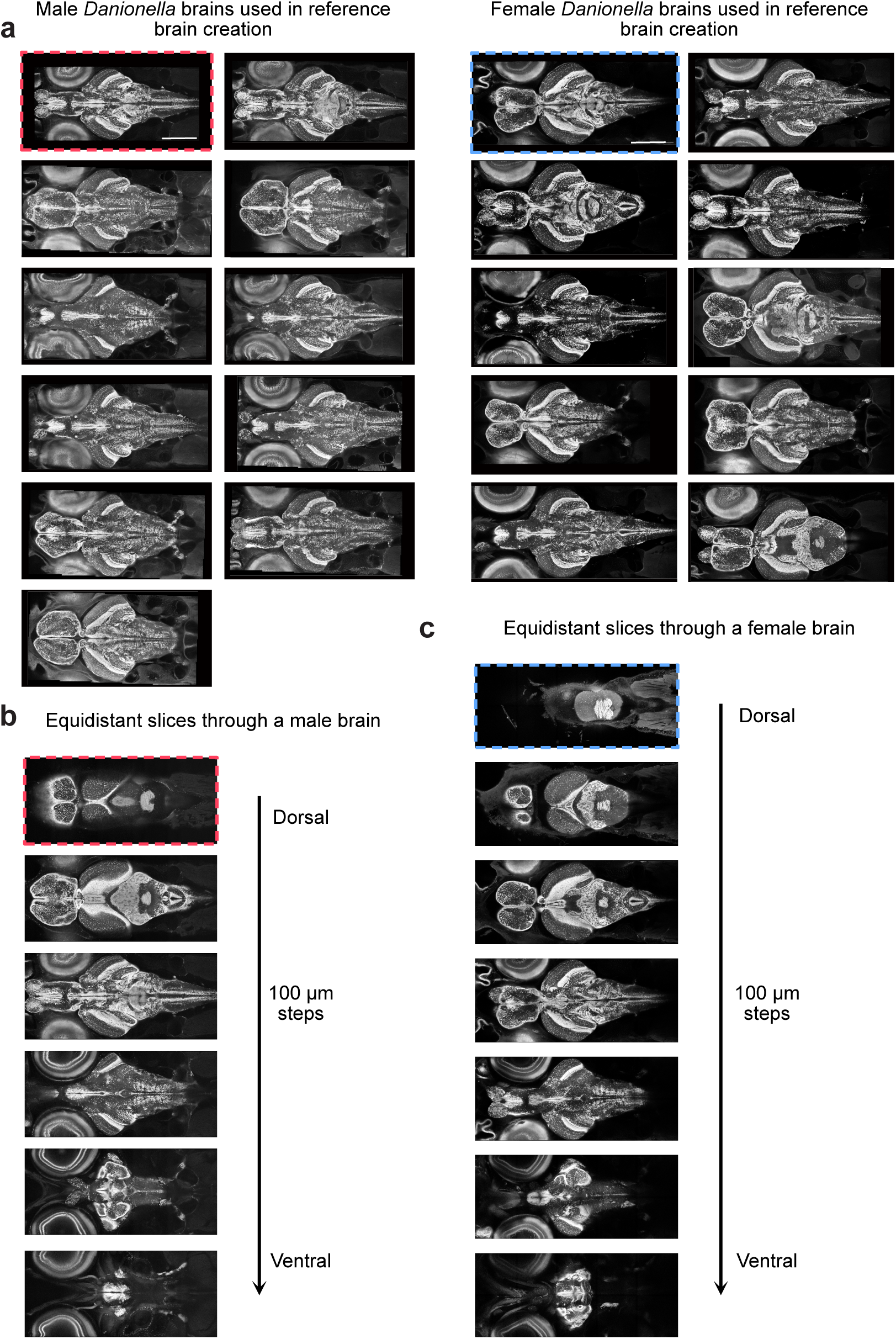
Anatomical brain stacks used in the creation of the reference brain. (a) 11 male (left two columns) and female (right two columns) brain volumes used in the reference creation. Each presented as a middle slice along the dorsoventral axis. Dashed outlines indicate brains used in (b) and (c). (b) Outlined male brain volume presented as slices every 100 μm through the entire volume along the dorsoventral axis to show cellular resolution throughout the entire volume. (c) Same as (b) for the outlined female brain. Scale bars, 500 μm.

**Figure S2.**
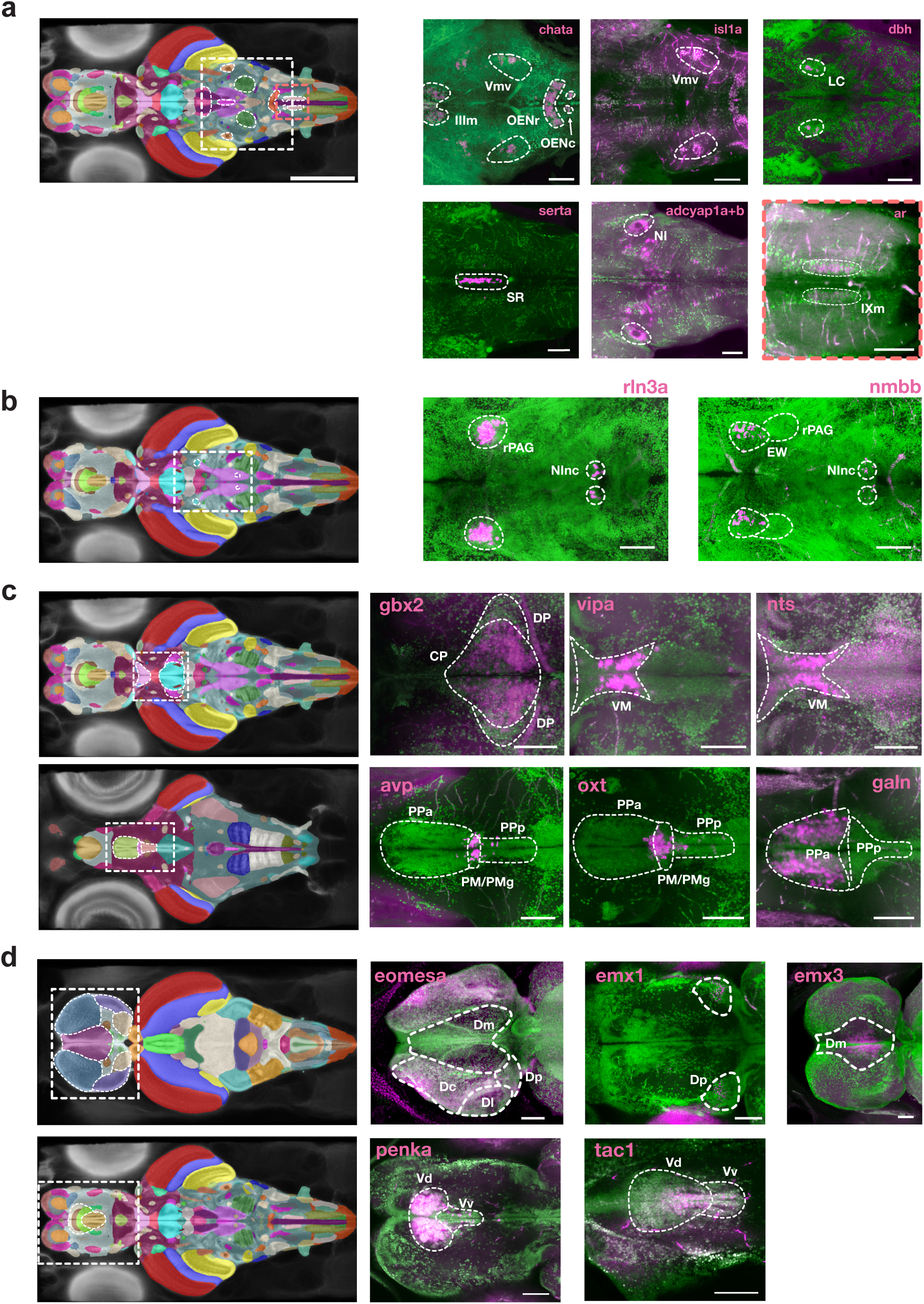
Full-resolution stainings used to support the anatomical segmentation of the *Danionella cerebrum* brain. Each image is a slice or a maximum intensity projection across the region of interest, with green showing the reference GCaMP channel expressed in nuclei and magenta showing a corresponding transcript staining using HCR 3.0. All volumes are presented before downsampling and registration to show the full expression pattern. The brain insets on the left show the approximate corresponding slice from the segmented atlas, with dashed lines outlining the shown fields of view and areas of interest that were annotated based on these stainings. Inset scale bars, 500 μm. (a) Annotated areas in the hindbrain using *chata* (midbrain oculomotor nucleus IIIm, trigeminal motor nucleus Vmv and rostral and caudal octavolateralis effect neurons OENr and OENc)*, isl1a* (highlighting trigeminal motor nucleus Vmv)*, dbh* (locus coeruleus LC)*, serta* (superior raphe SR)*, adcyap1* (mixed probes for *a* and *b* gene copies highlighting isthmic nucleus NI) and *ar* (highlighting weakly stained motor nucleus of the ninth cranial (glossopharyngeal) nerve IXm). (b) Rostral and caudal extent of the periaqueductal grey PAG / central grey CG in *Danionella cerebrum.* Rostral population is identified through the non-overlapping *rln3a* expression bordering the non-PAG Edinger-Westphal nucleus EW expressing *nmbb.* Both of these expression patterns overlap in the caudal PAG nucleus incertus NInc. (c) Annotation of the diencephalic thalamic and preoptic area divisions. Top row: *gbx2* identifies the dorsal thalamic nuclei (CP and DP). *vipa* and *nts* expression identify the rostrally located ventromedial nucleus VM. Bottom row: *avp* expressing populations mostly in the magnocellular preoptic nucleus PM with the gigantocellular part PMg divides the anterior PPa and posterior PPp parvocellular preoptic nuclei. *oxt* expression is partially overlapping with *avp* but extending ventrally and caudally. Finally, *galn* expression is restricted to the anterior PPa. (d) Annotations of the telencephalic divisions. Top row: pallial divisions are separated by *eomesa* (Dc, Dl, Dp, but not Dm), *emx1* (Dp) and *emx3* (Dm). Bottom row: dorsal division of the subpallial ventral telencephalon (Vd) express *penka* and *tac1,* separating it from the ventral nucleus of the ventral telencephalon Vv. Staining scale bars, 100 μm.

**Figure S3.**
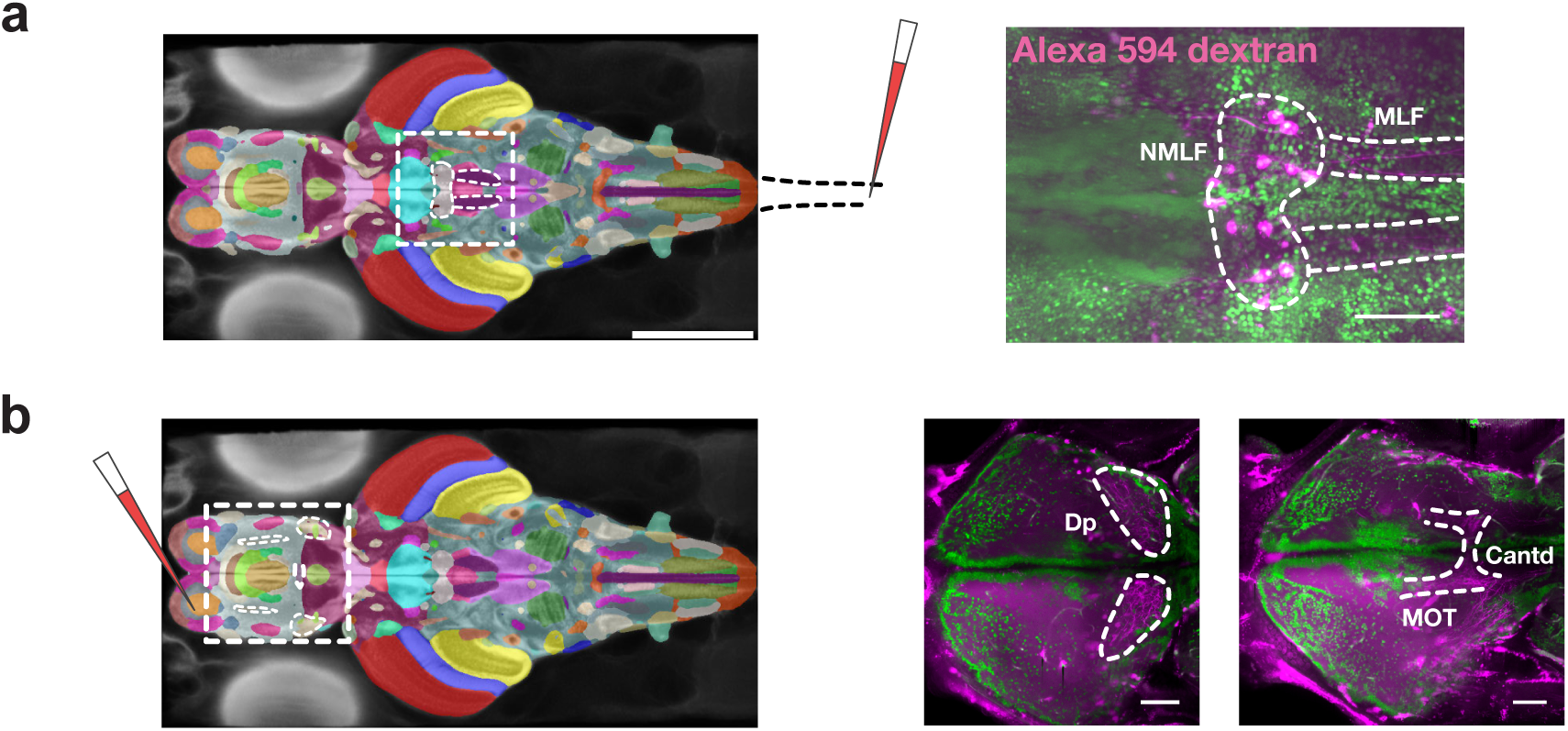
Tracer injections of the Alexa 594 dextran. (a) Rostral extent of the spinal backfill confirms the position of the nucleus of the medial longitudinal fascicle NMLF, with individual processes visible within the MLF. (b) Olfactory bulb injection tracing fibers through the telencephalon to confirm the location of the olfactory Dp (left). Individual fibers in the medial olfactory tract MOT visibly crossing the dorsal anterior commissure Cantd (right). Scale bars, 100 μm.

**Figure S4.**
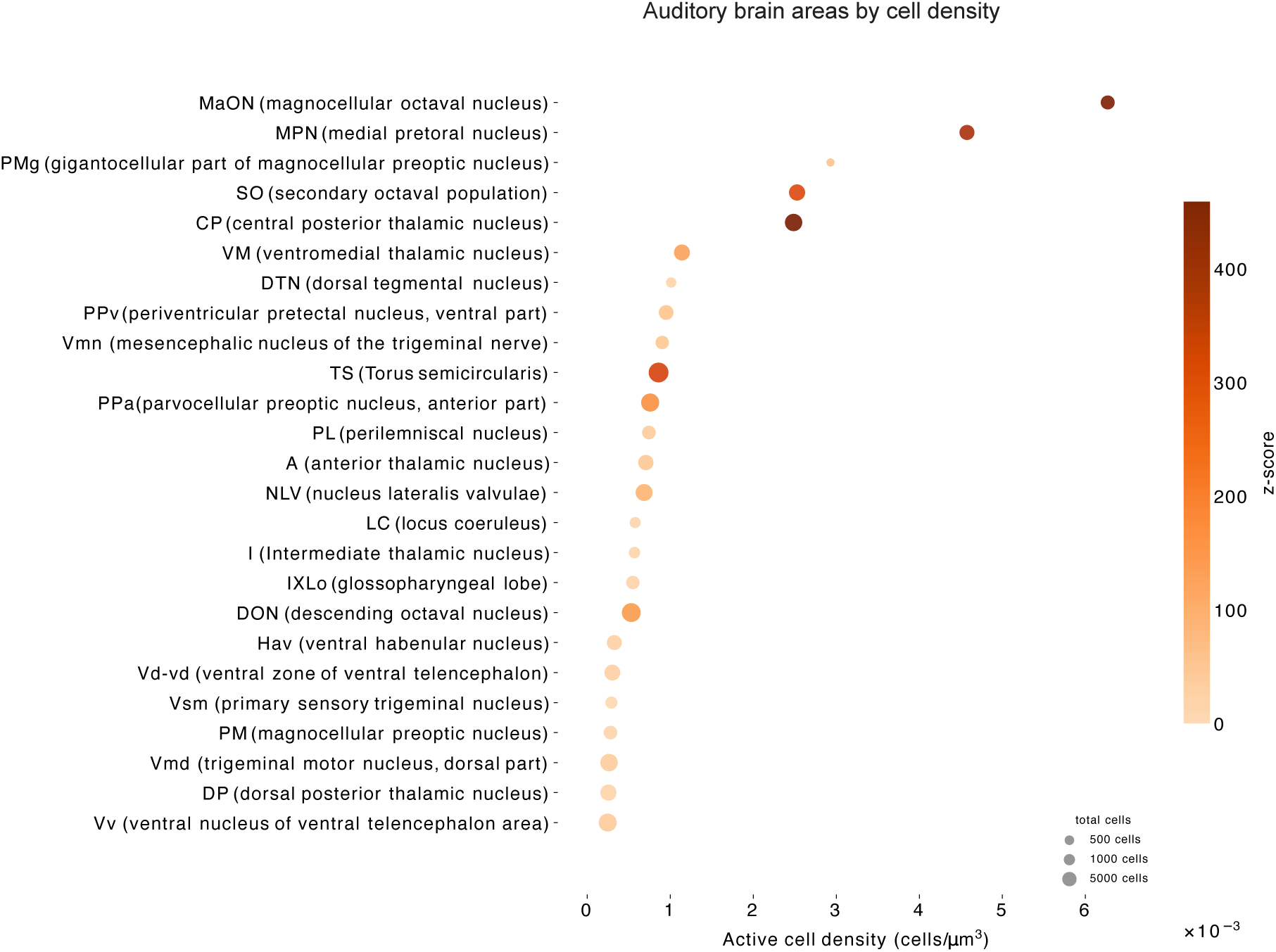
Areas with highest density of auditory-responsive cells. Top 25 areas with significantly overrepresented auditory stimulus-responsive cells sorted by responsive cell density. Significance: for all, p < 0.05 permutation testing with per-fish stratification (see Methods). Z-score indicates the deviation of the observed cell distribution from the permutation-based null distribution.

## Supplementary tables

The following tables are attached separately:

**Supplementary Table 1.** List of all target sequences used for the design of the HCR 3.0 staining probes and amplifiers.

**Supplementary Table 2.** Table of all annotated brain areas with their ID, abbreviation, full name and a note supporting their identification, with references where applicable. All areas without a note are annotated based on direct comparison with the AZBA zebrafish brain segmentation.

